# Injection practices in 2011-2015: A rapid review using data from the Demographic and Health Surveys (DHS)

**DOI:** 10.1101/574137

**Authors:** Tomoyuki Hayashi, Yvan J-F Hutin, Marc Bulterys, Arshad Altaf, Benedetta Allegranzi

## Abstract

**Background:** Reuse of injection devices to give healthcare injections decreased from 39.8% to 5.5% between 2000 and 2010, but trends since 2011 have not been described. We reviewed results of Demographic and Health Surveys (DHS) to describe injection practices worldwide from 2011 to 2015.

**Methods:** We searched the DHS Internet site for data published on injection practices conducted in countries from 2011 to 2015, extracted information on frequency (number of healthcare injections per person in the last 12 months) and safety (proportion of syringes and needles taken from a new, unopened package). We compared gender groups and WHO regions in terms of frequency and safety. For countries with data available, we compared injection practices 2004- 2010 and 2011-2015.

**Results:** Since 2011, 40 of 92 countries (43%) that had DHS surveys reported on injection practices. On average, the frequency of injection was 1.64 per person per year (from 3.84 in WHO Eastern Mediterranean region to 1.18 in WHO African region). Among those, 96.1% of injections reportedly used new injection devices (from 90.2% in the WHO Eastern Mediterranean region to 98.8% in the WHO Western Pacific region). On average, women received more injections per year (1.85) than men (1.41). Among 16 (40%) countries with data up to 2010 and since 2011, 69% improved in terms of safety. The annual number of unsafe injections was reduced in 81% of countries, with the notable exception of Pakistan where the number of unsafe injections was the highest and did not decrease between 2006 and 2012.

**Conclusion:** Injection practices have continued to improve in most countries worldwide, although the Eastern Mediterranean region in particular is facing residual unsafe practices that are not improving. Further efforts are needed to completely eliminate unsafe injection practices in health care settings, including through the use of reuse-prevention devices. Despite some limitations, DHS is an easily available method to measure progress over time.

## Introduction

A safe injection never harms the recipient, does not expose the provider to avoidable risks and does not result in any waste that is dangerous to other people.^1^ The World Health Organization (WHO) estimated that in 2000, 16 billion healthcare injections were given each year in developing and transitional countries. ^2^ Of these, 90-95% were for therapeutic purposes, while 5-10% were immunizations.^2^ Injections are often used unnecessarily when oral medicine could be equally effective.^3^ WHO estimated that in 2000, 39.8% of injections were given with devices reused in the absence of sterilization. ^2^ Health care injections given with re-used equipment expose patients to infections with bloodborne pathogens, including hepatitis B virus (HBV),^4^ hepatitis C virus (HCV),^5^ and human immunodeficiency virus (HIV).^6^ Overuse of injections to administer medicines amplifies the risk of transmission. WHO estimated that in 2000, overuse and unsafe use of healthcare injections caused 30% of new infections with HBV (21 million), 41% of new infections with HCV (2 million), and 9% of new infections with HIV (260,000, annually).^7^

Since 2000, WHO worked on the establishment of policies for the safe and appropriate use of injections worldwide. The Safe Injection Global Network (SIGN) regrouped efforts from all stakeholders, including international organizations, governments, nongovernmental organization, civil society, and industry.^8^ From 2004, the Demographic and Health Surveys (DHS) started to include data on injection practices for surveys conducted in several countries. ^9^ DHS are nationally representative population-based surveys of adult population with large sample sizes (e.g., > 5,000 households). The DHS Programme has provided technical assistance to more than 300 surveys in over 90 countries, advancing global understanding of health and population trends. New questionnaire items included injection frequency (the average number of healthcare injections reported per person per year) and injection safety (whether for the last healthcare injection, the syringe and needle came from a new, unopened package). In the year 2010, WHO commissioned an update of the systematic review on injection practices, which largely used DHS data. The results indicated that between 2000 and 2010, the proportion of reuse of injection devices dropped from 39.8% to 5.5%.

Meanwhile, the frequency of use of injection to administer medications had not decreased.^10^ Overuse and unsafe use of injections still led to transmission of bloodborne pathogens, although less so than in 2000. From 2000 to 2010, despite a 13% population growth, there was an estimated reduction of 87% and 83%, respectively, in the absolute numbers of HIV and HCV infections transmitted through healthcare injections. For HBV, the reduction was more marked (91%) due to the additional impact of vaccination.^11^

In 2016, WHO published injection safety guidelines recommending safety engineered injection devices to eliminate unsafe injections.^12^ The WHO policy document specifically addressed the use of these devices for therapeutic injections. Further, WHO launched a comprehensive package of implementation tools and other resources to facilitate adoption of the injection safety recommendations according to a multimodal implementation strategy in 2017.^13^ More recent and higher quality data are needed to monitor the evolution of injection safety since 2011, including to track progress before and after the launch of the WHO policy. ^14^ However, systematic reviews can be costly and time consuming. The ongoing collection of injection practices information within DHS provided an opportunity to obtain new estimates easily and rapidly. The objectives were to describe injection practices since 2011, including heterogeneity by region and gender, and to determine if injection safety has improved since 2011.

## Materials and Methods

We collected and reviewed healthcare injection practices worldwide in terms of frequency and safety according to the indicators conducted in DHS between 2011 and 2015 (since 2011). If countries surveyed since 2011 had a prior DHS survey data including injection practice up to 2010, we included the prior survey to compare data up to 2010 and since 2011.

### Data collection during the DHS surveys

DHS questionnaires were conducted in countries and adapted from template survey instruments developed by the DHS programme to reflect the population and health issues relevant to each country. The questionnaire was translated into each native language. A nationally representative sample of households (women age 15-49 and men age 15-49, 54, or 59) was eligible for individual interviews that were performed individually. We focused on two template questions. These were ‘have you had an injection for any reason in the last 12 months? If yes, how many injections have you had?’ and ‘the last time you got an injection from a health worker, did he/she take the syringe and needle from a new, unopened package?’. When considering DHS questionnaire items, we used the term ‘frequency’ to refer to the average number of health care injections per person in the last 12 months, and used the term ‘safety’ to refer to the proportion of injections for which the respondent mentioned the syringe and needle had been taken from a new, unopened package.

### Search for DHS surveys

We accessed the homepage of DHS Program (https://dhsprogram.com/) and searched all DHS final reports in 2011-2015 for data on injection practices. If the surveyed year spanned over two years (such as 2015-16), we considered the survey to have been conducted in the year it was initiated. When the data was available, we also searched the survey data conducted up to 2010 in the same country for comparison. The data was available in 2004-2010.

### Data abstraction

We extracted the data on frequency and safety. The denominator for frequency was the total number of respondents. The denominator for safety was the number of respondents who had received at least one health care injection in the last 12 months.

### Data analysis

Some countries had only information on injections received in the last six months. In that case, we doubled frequency to adjust for the different referent period. We combined the data on frequency and safety to calculate the number of unsafe injection per person per year [(the average number of health care injections per person in the last 12 months) x {1-(the proportion of injections from a new package)}]. For countries that had surveys conducted at the subnational level, we calculated the average value of the subnational surveys.

We combined the data from the various surveys to estimate the global situation in terms of frequency and safety. We compared gender groups, regions, and countries in terms of the number of unsafe injections. For countries that had more than two DHS surveys since 2004, we compared practices up to 2010 and since 2011 in terms of safety and frequency. If there were multiple surveys up to 2010, we adopted the most recent one to reflect on the most recent evolution. For countries that only had data on women, we restricted the comparison to women up to 2010 and since 2011.

## Results

### DHS surveys available

We identified 92 countries with DHS data available between 2004 and 2015 as of December 2017. Among these countries, 43% (40/92 countries) included injection safety data in 2011-2015 (Table 1). Among these 40 countries, 40% (16/40 countries) also had the data up to 2010 available for comparison over time (Table 1).

**Table 1:** 40 countries with injection practices data -in DHS surveys conducted between 2011-2015 and 16 countries with corresponding prior data (2004-2010).

| Region | Country | 2011-2015 | 2004-2010 |
| --- | --- | --- | --- |
| African | Benin | 2011 | 2006** |
| African | Cameroon | 2011 | No data |
| African | Congo | 2011 | No data |
| African | Cote d'Ivoire | 2011 | No data |
| African | Equatorial Guinea | 2011 | No data |
| African | Ethiopia | 2011 | 2005 |
| African | Mozambique | 2011 | No data |
| African | Uganda | 2011 | 2006 |
| African | Comoros | 2012 | No data |
| African | Gabon | 2012 | No data |
| African | Mali | 2012 | 2006** |
| African | Niger | 2012 | No data |
| African | Gambia | 2013 | No data |
| African | Liberia | 2013 | 2007 |
| African | Namibia | 2013 | 2006 |
| African | Nigeria | 2013 | 2008 |
| African | Sierra Leone | 2013 | 2008 |
| African | Togo | 2013 | No data |
| African | Zambia | 2013 | 2007 |
| African | Chad | 2014 | No data |
| African | Ghana | 2014 | 2008 |
| African | Kenya | 2014 | No data |
| African | Lesotho | 2014 | No data |
| African | Rwanda | 2014 | 2005, 2010 |
| African | Tanzania | 2015 | 2004, 2010** |
| African | Zimbabwe | 2015 | 2005, 2010 |
| Americas | Honduras | 2011 | No data |
| Americas | Haiti | 2012 | 2005 |
| Americas | Dominican Republic | 2013 | No data |
| Eastern Mediterranean | Pakistan | 2012 | 2006** |
| Eastern Mediterranean | Yemen | 2013 | No data |
| Eastern Mediterranean | Afghanistan | 2015 | No data |
| European | Kyrgyz Republic | 2012 | No data |
| European | Tajikistan | 2012 | No data |
| European | Armenia | 2015 | 2010 |
| South-East Asia | Nepal | 2011 | No data |
| South-East Asia | Indonesia | 2012 | No data |
| South-East Asia | India* | 2015 | No data |
| South-East Asia | Myanmar | 2015 | No data |
| Western Pacific | Cambodia | 2014 | No data |

\* Average value of the five areas surveyed
\*\* Use only female data in comparison because no male data available

### Safety and frequency

Overall, respondents reported 1.64 injections per person per year (N= 840,711) in 2011-2015. Of these, 96.1% (N= 279,620) were reported to have been given with devices taken out of a new packet (i.e. were considered to be safe).

### Regional variations

The Eastern Mediterranean Region (EMR) had the highest number of unsafe injections (0.38 per person per year). EMR also had the highest average number of health care injections per person in the last 12 months and the lowest proportion of injections with devices from a new package (Table 2 and Figure 1). EMR was followed by the South-East Asia Region (SEAR, 0.18 per person per year), the European Region (EUR, 0.08 per person per year), the African Region (AFR, 0.04 per person per year), the region of the Americas (AMR, 0.02 per person per year), and the Western Pacific Region (WPR, 0.02 per person per year).

**Table 2:** Injection practices in WHO regions according to DHS surveys conducted in 2011-2015 Figure 1: Annual number of safe and unsafe injections, by WHO region, DHS surveys conducted in 2011-2015

| Region | Frequency |  | Safety |  |
| --- | --- | --- | --- | --- |
|  | Health care injections per person per year | Sample size | Use of an unopened syringe or needle | Sample size |
| **African** | 1.18 | 466,836 | 96.7% | 144,115 |
| **Americas** | 1.28 | 73,335 | 98.4% | 12,951 |
| **Eastern Mediterranean** | 3.84 | 82,347 | 90.2% | 32,467 |
| **European** | 2.92 | 29,148 | 97.1% | 6,881 |
| **South-East Asia** | 2.66 | 166,277 | 93.3% | 75,233 |
| **Western Pacific** | 1.60 | 22,768 | 98.8% | 7,973 |
| **Total** | 1.64 | 840,711 | 96.1% | 279,620 |

**Figure 1:**
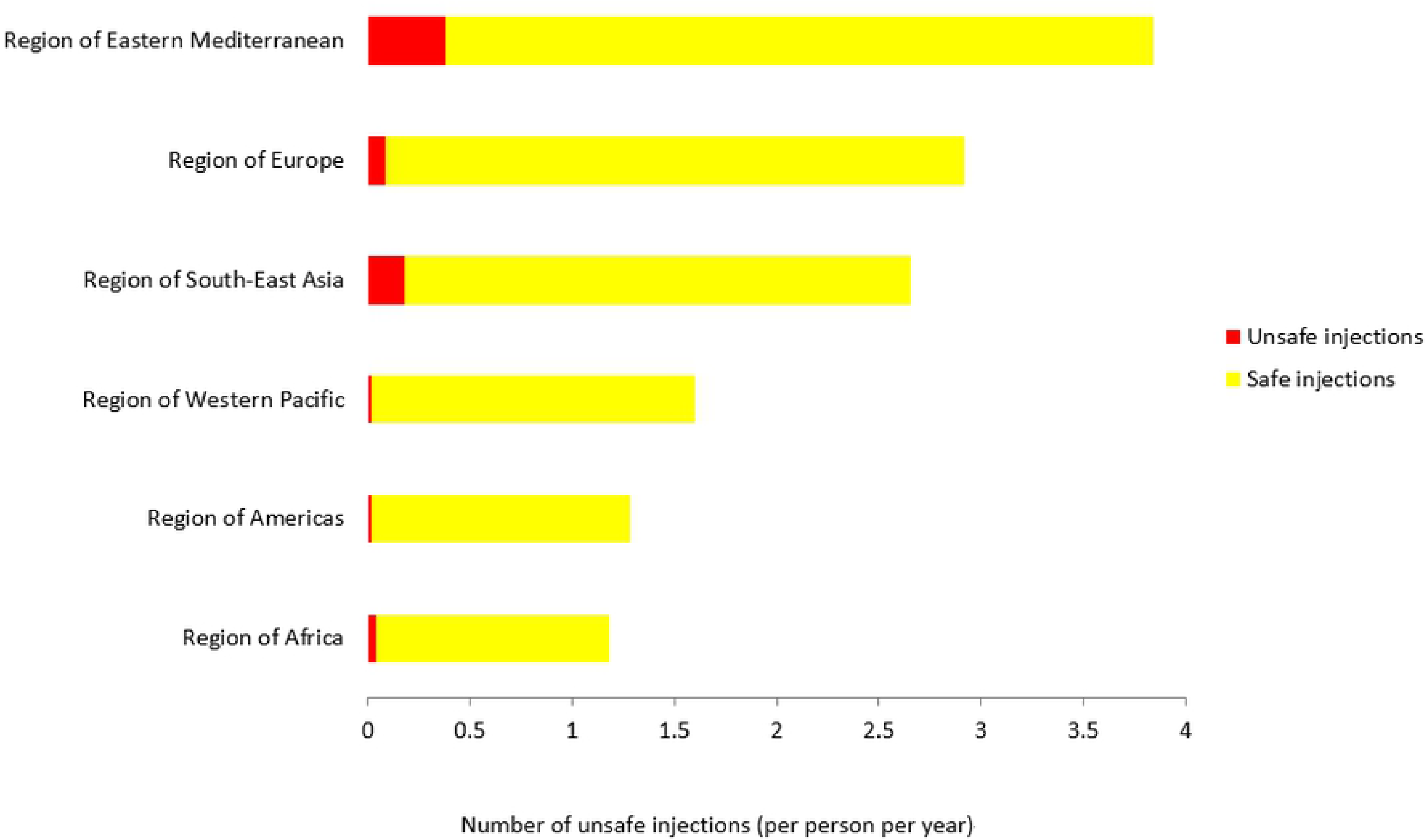
Annual number of safe and unsafe injections, by WHO region, DHS surveys conducted in 2011-2015

### Analysis by gender

The average number of health care injections per person in the last 12 months was higher in women (1.85 per person per year, sample size: 575,490) than in men (1.41 per person per year, sample size: 265,221). However, the proportion of injection devices taken from a new package was almost the same in women (96.2%, sample size: 202,969) and in men (96.0%, sample size: 76,651). Overall, the number of unsafe injections per person and per year was 0.071 in women and 0.056 in men.

### Evolution over time

Among the 16 countries with data up to 2010 and since 2011, the average annual number of injections decreased from 1.44 (sample size: 252,513) to 1.25 (sample size: 317,886). Similarly, the average proportion of safe injection increased from 95.9% (sample size: 70,953) to 97.0% (sample size: 98,103, Table 3). In terms of frequency, 9 of 16 countries (56%) had improved (average improvement: 0.49, range: 0.10-2.35) and 9 of 16 (38%) had deteriorated (average deterioration: 0.18, range: 0.05-0.30). In terms of safety, 11/16 countries (69%) improved (average improvement: 1.7%, range: 0.2-5.1%) and 5/16 countries (31%) deteriorated (average deterioration: 0.8%, range: 0.1-1.4%). The annual number of unsafe injections improved in 81% (13/16 countries) and deteriorated in 19% (3/16 countries). In Pakistan, where the number of unsafe injection was the highest among the 16 countries, the number of unsafe injection did not change substantially between 2006 (0.71 per person per year) and 2012 (0.80 per person per year).

**Table 3:** Comparison of injections practices up to 2010 and since 2011 in the 16 countries where DHS data were available for comparison

|  | Frequency |  | Safety |  |
| --- | --- | --- | --- | --- |
|  | Health care injections per person per year | Sample size | Use an unopened syringe or needle | Sample size |
| **Up to 2010\*** | 1.44 | 252,513 | 95.9% | 70,953 |
| **Since 2011** | 1.25 | 317,866 | 97.0% | 98,103 |

\* Use only more recent data if there are multiple data.

## Discussion

In 2010, a review of injection practices globally reported improvement in injection safety following the implementation of joint efforts from WHO, partners of the Safe Injection Global Network, other agencies and countries.^10,11^ However, it also pointed to the absence of improvement in reducing the frequency of health care injections annually. This new rapid review provides an indication of the evolution of the situation since 2011, and sheds light on the additional progress and setbacks.

The most recent DHS surveys suggested that in 2011-2015, the trend towards improvement of injection practice continued. However, injection frequency and safety continued to vary widely by region. The WHO Eastern Mediterranean region has the highest number of injections and the highest proportion of unsafe injections, followed by the WHO South-East Asia Region (SEAR). The number of unsafe healthcare injection in the WHO Eastern Mediterranean Region (EMR) was also the highest in the world in 2000 (2.96 per person per year)^2^ and in 2010 (0.57 per person per year).^10^ In these regions, a high proportion of injections are administered by private providers who may have no formal medical qualification. In such informal settings, health care providers’ attitudes also drive injection overuse. The reference to standard treatment guidelines is uncommon. Injections are frequently used on an “ad hoc” basis to administer mixtures of antibiotics, analgesics, vitamins or anti-histamines in the desire to meet user demand.^15,16^ Reducing injection overuse would only be a matter of promoting rational use of medicines if injections were administered safely.^17^ However, reuse of syringes frequently leads to transmission of bloodborne pathogens.^18,19,20,21,22,23,24,25^ In EMR and SEAR, especially India and Pakistan, blood-borne pathogens are prevalent and can be transmitted, around 1.7%-5.6% for HBV,^26^ 0.5%-3.8% for HCV,^27^ and 0.3%-0.1% for HIV.^28^ For instance, unsafe health care injections remain a key driver of the HCV epidemic in Pakistan.^29^ This could be prevented as patients’ knowledge regarding transmission of bloodborne pathogens drives consumer demand for new syringes.^30,31^

According to DHS, women generally received more injections than men. The reasons for this may include obstetrical and gynecological conditions, including contraception, delivery, and prevention of maternal and neonatal tetanus. DHS data also suggest that there is no difference in injection safety between women and men. Some of the injections received by women could be received through programmes such as immunization or contraceptive treatment rather than informal health care. However, the absolute number of unsafe injections was higher in women than in men. Actually, exposure to healthcare (e.g. health care injections, hospitalizations and pregnancies) has been associated with HCV infection among women in Pakistan.^32,33^ Addressing this issue has the potential to reduce gender inequalities in injection safety. Since 1999, the principle of “bundling” has been proposed to ensure the safety of injections given through key public health programmes, so that the sources of financing of immunization and contraception commodities also pay for safe injection devices and safety boxes for disposal of used injection equipment.^34^ Bundling refers to the concept of a theoretical “bundle” which comprises each of the following items: 1) good quality vaccines, 2) auto-disable syringes, and 3) safety boxes. “Bundling” has no physical connotation and does not imply that items must be “packaged” together.^35^

Comparison of the data up to 2010 and since 2011 in 16 countries with available data suggested an improvement in many countries. Concerted efforts of ministries of health, local health facilities, non-profit organizations, and donors may explain some of the progress. However, in Pakistan, Uganda and Zambia, the number of unsafe injections did not change substantially since 2011. Pakistan is a country with a rapidly increasing population. The effects of the reduction in unsafe injections might be reduced by the population growth in some densely populated regions.^36^ However, in theory, the scale up of safe injections could be more cost effective and positively associated with infections averted in growing populations.^37^ This is important because Pakistan has the second highest number of HCV infections in the world.^27^

Injection-associated transmission of bloodborne pathogens can be prevented through the development of a strategy to reduce injection overuse and achieve injection safety and its implementation by a national coalition.^38^, The achievement of this critically important goal requires the establishment of a national multidisciplinary coalition involving different departments of the Ministry of Health and other stakeholders, such as non-governmental organizations and private health-care providers. A behaviour change strategy targeting consumers as well as the public, private and lay health-care workers should be strongly advocated. Equipment and supplies should be provided to eradicate the re-use of syringes and needles without sterilization. Sharps waste should be managed to ensure that disposable syringes and needles are not re-used and do not lead to needlestick injuries.^38^ In addition, since 2016, WHO recommends a policy to prefer the use of sharp injury protection syringes for intramuscular, intradermal and subcutaneous injections over the use of re-use prevention syringes for intramuscular, intradermal and subcutaneous injections.^12^ Syringes with sharps injury prevention features reduce the incidence of needlestick injuries.^39^ Healthcare workers perceive injection safety devices to be generally easy to use, safe, and tolerated by patients.^40^ Rapid implementation of this policy could address residual injection safety issues in resource-limited settings where unsafe practices persist.

Our review and analysis of existing data have a number of limitations. Some limitations are inherent to the methods used by DHS. Survey respondents might not notice or remember whether the injection they are receiving was given with a new needle and syringe from an unopened package when interviewed months later (recall bias). Furthermore, the respondents might tend to answer questions in a manner that will be viewed socially favorable because the data was collected by questionnaire (social desirability bias). There are also limitations due to our rapid review methods. First, we did not calculate confidence intervals, did not perform statistical comparisons and did not model regional or global estimates. Calculation of confidence intervals in statistical analyses is based on the assumption of random sampling, yet the countries that were included in the DHS programme were not chosen at random. Further, the low proportion of unsafe injections, the lack of variability in terms of income groups and the limited sample size of countries included would have limited efforts to model regional and global estimates. Second, we used only one data source (DHS), which provides limited evidence in some areas while there could be other data sources for injection safety. Third, doubling the six-month recall may overestimate the response that would’ve been generated by asking people to recall 12 months. Fourth, we aggregated estimates from countries that had different samples of their population with respect to different age group and gender. Finally, there is large variation globally by region and a lack of data from many countries. Overall, our estimation is an imperfect reflection of the current status worldwide. The review is limited to healthcare injections, whereas in most countries non-healthcare injections, including injecting drug use, play an important role in the transmission of bloodborne infections through unsafe injections.

In conclusion, despite substantial improvements in the past decade, unsafe injections persist worldwide, and continue to vary widely by WHO regions. Women continue to be at higher risk for the consequences of unsafe injections than men. Reported injection practices continued to improve in many countries, but improvements appear to have levelled off since 2011 in some countries, and the situation may have deteriorated in others. On the basis of these findings, we suggest a number of actions. First, strong promotion of the safe and appropriate use of injections is needed, particularly in regions most affected by unsafe injection practices, including the WHO Eastern Mediterranean and South-East Asia regions. Use of WHO recommendations and implementation tools can facilitate implementation.^12^ Second, we must address injection safety in the field of reproductive and maternal health services to eliminate any gender inequalities concerning injection safety. Third, stringent efforts are needed to prevent injection-associated infections with bloodborne pathogens through improved technologies. A targeted approach with dissemination of auto-disable and re-use prevention syringes in resource-limited settings where safety remains an issue and the risk of injection-associated infection is high could eliminate unsafe injections. Finally, there is a need to collect high-quality data on injection safety worldwide and to make use of the available DHS data and other sources of information to justify, guide, and evaluate national safe and appropriate use of injection policies.

## Acknowledgement

The authors wish to acknowledge the comments of Dr Steve Luby on this manuscript.

## Disclaimer

The authors alone are responsible for the views expressed in this article and they do not necessarily represent the views, decisions or policies of the institutions with which they are affiliated.

